# Synergy of LPS and Shiga-toxin exposure on the RNA expression program of cultured human umbilical vein endothelial cells

**DOI:** 10.1101/2024.10.30.621129

**Authors:** David Svilar, Martijn A. van der Ent, David Siemieniak, David Ginsburg, Audrey C. Cleuren

## Abstract

**Background:** Hemolytic uremic syndrome (HUS) is a primary thrombotic microangiopathy that can cause acute renal failure, particularly in children, and is commonly triggered by Shiga toxin-producing *Escherichia coli* (STEC) infection. Although previous *in vitro* studies have shown that Shiga toxin (STX) can directly affect the endothelium, gram-negative bacteria such as *E. coli* also contain lipopolysaccharide (LPS) which is known to independently activate endothelial cells. Furthermore, studies in mice and baboons have shown a synergistic effect of LPS and STX in inducing HUS when administered simultaneously.

**Objective:** Determine how LPS, STX, or combined LPS/STX treatment affect the gene expression program of endothelial cells.

**Methods:** Human umbilical vein endothelial cells (HUVECs) were treated with STX, LPS or a combination of STX and LPS for either 4 or 24 hours, after which unbiased RNA sequencing analysis was performed.

**Results:** Four-hour treatment with LPS resulted in a greater number of differentially expressed genes as compared to treatment with STX. In contrast, 24-hour treatment with LPS or STX induced similar numbers of differentially expressed genes. Dual treatment with both toxins resulted in a synergistic effect, showing significant up-or downregulation of multiple transcripts that were not observed in either single treatment group, particularly at the 24-hour timepoint.

**Conclusion:** Our data provide a resource for assessing the role of endothelial cell gene expression in the pathogenesis of STEC infection and suggest that the responses of these cells to LPS, STX, or combined LPS/STX treatment are unique and temporally regulated.

## Introduction

Hemolytic Uremic Syndrome (HUS) is predominantly caused by infection with Shiga-toxin (STX) producing *Escherichia coli* (STEC) O157:H7. It is the most common primary thrombotic microangiopathy (TMA) worldwide, with the majority of patients developing acute renal failure (1-3). STX-induced damage is thought to activate microvascular endothelial cells (ECs), thereby promoting thrombotic microvascular injury in organs such as the gut, brain, heart, and kidney, with renal histopathology demonstrating glomerular capillary EC thickening, swelling and luminal thrombosis (3). STX can bind circulating white blood cells, which is thought to facilitate the transfer of active STX from the gut to kidney endothelial cells (4). Here, STX binds EC glycosphingolipid globotriaosylceramide (Gb3) and is retrograde transported through the Golgi and endoplasmic reticulum (ER), subsequently disrupting protein synthesis leading to endothelial cell injury (5). Although previous studies have confirmed a key role for Gb3 receptor synthesis in STX entrance and cytotoxicity (6, 7), how STX-induced EC injury promotes HUS is still unknown.

In addition to STX, STEC also produces other toxins, including lipopolysaccharide (LPS), which is known to cause many transcriptional changes independent of STX (8). Interestingly, previous studies have shown that human umbilical vein endothelial cell (HUVEC) survival is decreased during combined LPS and STX treatment (9, 10). *In vivo*, LPS and STX have also been reported to have a greater than additive effect on worsening kidney function, both in mouse and baboon models of HUS (11, 12), with dual LPS/STX treatment of mice resulting in a greater number of significantly altered transcripts in the kidney as compared to individual LPS or STX treatment (11). However, as this latter study evaluated changes in gene expression at the organ level, it is not possible to discern whether those changes occurred within the endothelial compartment specifically. We now report a comprehensive RNA sequencing analysis of the effects of LPS and STX (either alone or in combination) in HUVECs, to mimic the early stages of STEC infection *in vivo*. We show that LPS-induced changes occur more rapidly than those induced by STX, with combined LPS/STX treatment resulting in greater than additive effects on the EC gene expression program as compared to LPS or STX alone.

## Methods

### Cell culture

Primary pooled HUVECs (Lonza #C2519A) were cultured in supplemented endothelial growth media-2 (Lonza EGM-2 BulletKit, #CC-3162) per the manufacturer’s instructions. Holotoxin variant Shiga-toxin 2a (STX2; Phoenix Laboratory at Tufts Medical Center) was dissolved in phosphate buffered saline (PBS, Gibco #10010-023) at a stock concentration of 100 ng/µL and frozen at −80°C in single use vials that were thawed immediately prior to treatment. LPS from *Escherichia coli* 026:B6 was obtained from Sigma (L8274) and dissolved in PBS at a concentration of 500 µg/mL. HUVECs were seeded (passage 2; 50,000 cells/well) in 24-well plates 24 hours prior to treatment with PBS, STX (50 or 5 ng/mL), LPS (2.5 μg/mL), or combined STX and LPS.

### RNA isolation and sequencing analysis

After 4 or 24 hours of treatment, cells were washed with ice cold PBS followed by RNA isolation (Takara NucleoSpin RNA) and cDNA conversion using the SMART-Seq v4 Ultra Low Input RNA Kit (Takara, #634892). cDNA amplification, purification, shearing, and quality assessment were completed as previously described (13). Subsequent end repair, adapter ligation and amplification of barcoded cDNA was performed using the KAPA HyperPrep Kit (KK8504) combined with NEXTFLEX-96 DNA Barcodes (NOVA-514105). Quality of sequencing libraries was assessed using an Agilent 2100 Bioanalyzer, and libraries were sequenced as 100 paired-end sequencing reads on the NovaSeq6000. Sequence mapping to the transcriptome (GRCh37 human version 32) and quantification was performed using Salmon (14), with the resulting output being directly loaded into DESeq2 using tximport for downstream normalization and differential expression analysis (15, 16). An average raw read count of 20 per sample was used to filter out low abundant transcripts to reduce background noise prior to differential expression analysis. Of note, transcripts below this filter threshold were predominantly non-protein coding (83.8%), whereas transcripts above the filter threshold were primarily protein coding (87.1%), suggesting the filter was effective at removing background noise while capturing significant changes in gene expression. A false discovery rate (FDR or q-value; (17)) of <0.05 in combination with a log_2_ fold change <−1 or >1 was used to define statistically significant differential gene expression. Gene Ontology (GO) analysis was performed on gene lists passing the statistical threshold using the “Search Tool for Retrieval of Interacting Genes/Proteins” (STRING) database (18). Data visualization was performed using R, and Venn diagrams were created online using http://bioinformatics.psb.ugent.be/webtools/Venn/.

### Reanalysis of STX treated HUVEC microarray data

Microarray data from a previous study where human dermal neonatal microvascular endothelial cells (HMVECs) were treated with STX ((19): GSE32710) was accessed through the Gene Expression Omnibus (20) and reanalyzed using the GEO2R webtool, where the same statistical cutoffs used for our RNA sequencing data were used.

## Data accessibility

All sequencing data is available through the GEO database (GSE285207).

## Results and Discussion

### STX- and LPS-induced gene expression changes exhibit significant overlap

RNA-seq data from HUVECs were highly reproducible between biological replicates (Fig 1A), independent of treatment group or timepoint. Approximately 20% of the total genes detected in PBS-treated control HUVECs (2804 out of 14245 genes) showed time-dependent differential gene expression when comparing expression levels at 4-hours versus 24-hour. Although significantly upregulated genes (933 / 2804) did not show enrichment for specific biological processes by GO analysis, significantly downregulated genes (1871 / 2804 genes) showed enrichment for GO terms focused on vascular development and proliferation. This is consistent with the expected senescence observed in primary cells during increased time in culture (21), and the phenotypic drift previously reported for primary ECs grown in culture after being removed from their *in vivo* microenvironment (13). Therefore, all subsequent analyses were performed using vehicle treated controls at the same timepoint as the reference to distinguish toxin-induced changes in gene expression.

**Figure 1:**
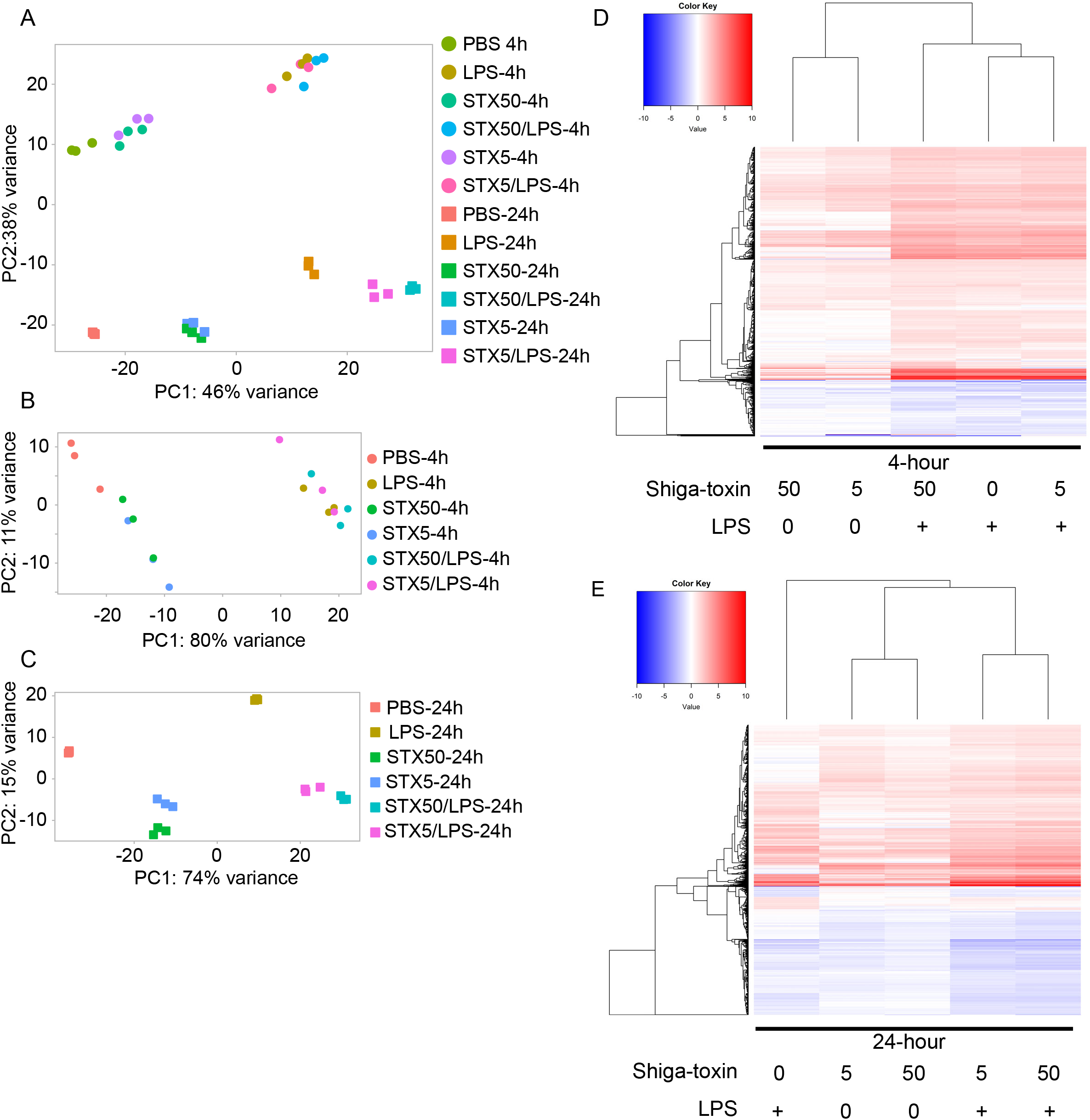
HUVECs treated with STX, LPS or dual treatment exhibit significant overlap of differentially expressed genes. Principal component analysis of vehicle, STX, LPS, and combined STX/LPS treatment at both (A) 4 and 24-hours, (B) 4-hours only, or (C) 24-hours only. Heatmap plotting the log_2_ fold change compared to vehicle controls, across treatments (columns) and individual genes (rows) for (D) 4-hours and (E) 24-hour treatment.

Upregulated genes composed the majority of the observed changes following toxin treatment at both time points (Figure 2A-F). Gene expression changes relative to PBS controls were similar for HUVEC treated with either 5 or 50 ng/ml STX at the 4-hour time point (Figure 1B). At 24-hours, a clear distinction between 5 or 50 ng/ml STX could be made, with STX 50ng/mL exhibiting a greater change in gene expression (Figure 1C). Therefore, we focused our analysis on this latter treatment group. Compared to PBS controls, LPS induced rapid changes in mRNA expression, with 1109 differentially expressed genes observed after 4-hour treatment, whereas only 309 genes were differentially expressed after STX treatment at 4 hours (Figure 2G). This is also reflected in the heatmap (Figure 1D), which indicates that the presence of LPS is the main driver of the group clustering. On the other hand, at 24 hours there were 1414 differentially expressed genes in the STX treatment group as compared to 1774 genes in LPS-treated cells, with 583 genes overlapping between the 2 treatment groups (Figure 2H) and STX treatment being the most evident driver of hierarchical clustering analysis (Figure 1E).

**Figure 2:**
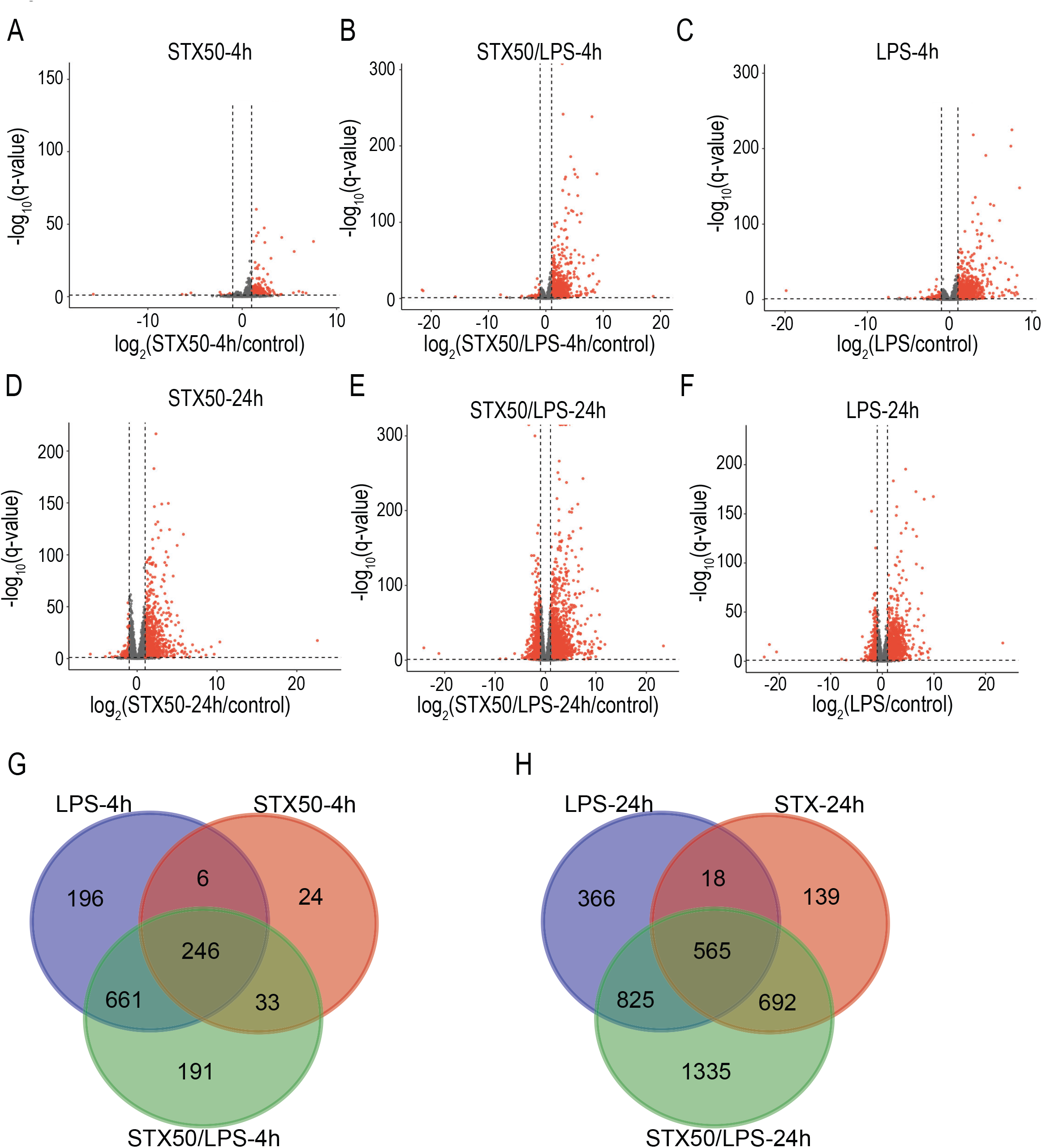
Volcano plots with differential expression plotted on the x-axis (log_2_ fold change of experimental condition compared to control), and statistical significance on the y-axis (−log_10_ of the q-value). Dashed lines demarcate the statistical cutoffs (log_2_ fold change <−1 or >1, q-value <0.05). Significantly differentially expressed genes (red), and genes not reaching the significance threshold (gray) were plotted for (A) STX 50 ng/mL at 4-hours, (B) STX 50ng/mL and LPS at 4-hours, (C) LPS at 4-hours, (D) STX 50ng/mL at 24-hours, (E) STX 50ng/mL and LPS at 24-hours, and (F) LPS at 24-hours. Venn diagrams display the overlap of differentially expressed genes between the different treatments compared to PBS controls at (G) 4-hours and 24-hours (H).

GO analysis of the 500 genes with the lowest q-values (17) after LPS treatment was conducted to assess enriched biological processes. As expected, LPS treatment resulted in differentially expressed genes associated with inflammatory processes and EC activation, which was sustained at the 24-hour timepoint. Interestingly, GO analysis of genes significantly affected by STX identified inflammatory processes limited to lymphocyte activation. For example, our data showed significant upregulation of TNF and IL6 with LPS treatment at both 4- and 24-hours, but not with STX treatment alone. As prior studies have shown that TNF and IL6 levels correlate with HUS severity and development of extrarenal complications (22), our results suggest that the LPS-induced inflammatory response in STEC infection likely directly contributes to HUS development.

STX treatment on the other hand, resulted in significantly upregulated transcripts overrepresented in GO terms focusing on the ER stress response and p38/MAPK signaling at 24 hours (Table 1). This is potentially due to STX-mediated protein synthesis inhibition inducing ER stress, consistent with previously reported findings that p38/MAPK signaling is essential for STX-induced increases in macrophage cytokine production (23, 24). ECs are known to increase cytokine production in response to bacterial toxins such as STX and LPS (25), which may also depend on p38/MAPK signaling. To assess whether these STX-induced changes were unique to HUVECs, we compared our data to a previously published study characterizing the response to 24-hour STX treatment in HMVECs, as evaluated by microarray analysis (19). The HMVECs used in the prior study were of dermal origin and composed of only microvascular ECs, thus differing from HUVECs in both the size of the vessel these ECs were isolated from as well as the tissue of origin. Although only 57 genes showed statistically significant differences in the HMVEC study, 35 of these genes overlap with our 24h STX treatment data. GO analysis of these 35 transcripts also identified enrichment for regulation of the p38/MAPK cascade, suggesting that this pathway is part of a general response to STX that is independent of EC origin.

**Table 1:** GO terms associated with significantly differentially expressed genes after 24-hour STX.

| GO term description | Strength | FDR | Proteins in network |
| --- | --- | --- | --- |
| Positive regulation of endoplasmic reticulum stress-induced intrinsic apoptotic signaling pathway | 1.21 | 0.0155 | SIRT1, PMAIP1, BBC3, DDIT3 |
| Histone H3 deacetylation | 1.14 | 0.0243 | SIRT1, PER1, SFPQ, HDAC9 |
| Inositol phosphate biosynthetic process | 1.1 | 0.0292 | ITPKC, IPPK, IPMK, ITPKB |
| Regulation of CD8-positive, alpha-beta T cell activation | 1.07 | 0.01 | IRF1, SH3RF1, RUNX1, CD274, CBFB |
| Nucleolus organization | 1.07 | 0.035 | RRN3, SIRT1, RRP8, USP36 |
| Integrated stress response signaling | 1 | 0.0016 | JUNB, FOS, EIF2AK3, FOSL1, ATF4, CEBPD, DDIT3 |
| Positive regulation of miRNA transcription | 0.92 | 0.00046 | NR3C1, EGR1, FOS, FOSL1, SMAD3, FOXO3, KLF4, BMP2, ETS1 |
| Adrenal gland development | 0.92 | 0.0324 | NR3C1, ARID5B, SMAD3, CITED2, CYP1B1 |
| Positive regulation of transcription by RNA polymerase I | 0.91 | 0.0121 | IPPK, BNC1, EIF2AK3, ATF4, DDX21, WDR43 |
| Regulation of p38MAPK cascade | 0.89 | 0.0052 | GADD45B, DUSP1, PER1, DUSP10, GADD45A, BMP2, CYLD |
| Regulation of endoplasmic reticulum stress-induced intrinsic apoptotic signaling pathway | 0.89 | 0.0137 | SIRT1, EIF2AK3, PMAIP1, PTPN1, BBC3, DDIT3 |
| Positive regulation of vascular endothelial growth factor production | 0.88 | 0.0407 | IL1A, EIF2AK3, ATF4, IL6, CYP1B1 |
| Positive regulation of response to endoplasmic reticulum stress | 0.86 | 0.0194 | SIRT1, PMAIP1, PTPN1, ERN1, BBC3, DDIT3 |
| Temperature homeostasis | 0.86 | 0.0496 | EGR1, NR1D1, IL1A, ADRB2, SQSTM1 |
| Placenta blood vessel development | 0.86 | 0.0496 | MAPK1, HES1, JUNB, FOSL1, HEY2 |

treatment, ranked by enrichment strength.

LPS facilitates STX binding to the cell surface by increasing Gb3 receptor expression in specific EC populations, including HUVECs (9). We observed a significant increase in both *UGCG* and *B4GALT5*, the 2 enzymes preceding the final enzyme to synthesize Gb3 (*A4GALT*), with LPS treatment. However, there was no significant difference in *A4GALT* expression between any of the treatment conditions. These data suggest the possibility that the previously reported increase in Gb3 in HUVECs after LPS treatment is due to differential expression of *UGCG* and *B4GALT5*, but not *A4GALT*, which may result in additional STX sensitivity with dual treatment. In this regard, renal ECs, a presumed primary STX target in HUS, have been reported to express 50x higher Gb3 at baseline and are more sensitive to STX than HUVECs, yet do not exhibit an increase in Gb3 or STX binding following LPS treatment (9). However, dual LPS/STX treatment has been shown to be synergistic in inducing HUS in both mouse (11, 26) and baboon models (12), thus suggesting that LPS treatment is conferring additional kidney EC sensitivity, despite significant Gb3 receptor present at baseline on renal ECs.

### Combined STX and LPS treatment have synergistic effects on gene expression

Since STEC produces both LPS and STX, we wanted to further test this potential interaction by treating cells simultaneously with both toxins. This led to significant expression changes in a unique subset of genes at 4 and 24 hours (Figure 2). Given the lower number of gene expression changes induced by STX at 4-hours, combined STX/LPS treatment had similar gene expression changes as compared to LPS treatment alone. This is also indicated by the overlap of data points in 4-hour PCA (Figure 1B), heatmap clustering (Figure 1D) and similar GO term enrichment, with 13 of the top 15 GO terms being identical. In contrast, of the total of 3940 genes differentially expressed at 24 hours following exposure to LPS, STX, or STX/LPS, 1335 were uniquely altered only in the combined treatment (Figure 2H). Pathway analysis of the 2057 genes upregulated after 24-hour dual STX/LPS treatment revealed enrichment of the unfolded protein response, p38/MAPK, NF-κB, and p-ERK signaling, whereas the 1360 transcripts downregulated by LPS/STX were associated with DNA repair and replication (Table 2). This latter finding suggests a decrease in cellular replication, which correlates with a prior report demonstrating that STX-treated cells slow their cell cycle progression (27).

**Table 2:** LPS/STX treatment for 24-hours decreased gene expression in GO terms related to.

| GO term description | strength | FDR | Proteins in network |
| --- | --- | --- | --- |
| Regulation of acyl-CoA biosynthetic process | 1.51 | 0.0107 | PDK4, PDK2, PDK3, SNCA |
| Positive regulation of DNA-directed DNA polymerase activity | 1.27 | 0.0016 | GINS2, CHTF18, DSCC1, PCNA, RFC4, RFC5 |
| Response to folic acid | 1.26 | 0.0363 | BCHE, MGMT, TYMS, CBS |
| Double-strand break repair via break-induced replication | 1.23 | 0.0107 | MCM5, GINS2, MCM2, CDC7, MCM3 |
| L-serine metabolic process | 1.23 | 0.0107 | SHMT1, PSAT1, PSPH, CBS, PHGDH |
| Serine family amino acid biosynthetic process | 1.13 | 0.0016 | SHMT1, CTH, PSAT1, PSPH, CBS, DHFR, PHGDH |
| Primary alcohol catabolic process | 1.1 | 0.0238 | SULT1E1, ALDH2, SULT1C4, AKR1C3, SULT1A1 |
| DNA unwinding involved in DNA replication | 1.04 | 0.0116 | MCM5, GINS2, RPA1, MCM2, MCM3, RAD51 |
| Nuclear DNA replication | 1.01 | 0.0158 | GMNN, MCM2, FEN1, PCNA, MCM3, RAD51 |
| DNA replication initiation | 0.99 | 0.0062 | CDC6, MCM5, MCM2, PRIM1, ORC1, CCNE2, MCM3 |
| Regulation of DNA-templated DNA replication | 0.97 | 1.11e-05 | MCM5, GMNN, E2F8, GINS2, CHTF18, MCM2, DSCC1, PCNA, RFC4, CDC7, RFC5, TIMELESS, MCM3 |
| Basement membrane organization | 0.91 | 0.0363 | MMP11, NID2, NID1, SPINT2, COL3A1, NTN4 |
| DNA duplex unwinding | 0.77 | 0.00058 | MCM5, GINS2, RPA1, BRIP1, CHTF18, MCM2, DSCC1, ERCC6L, RAD54L, RFC4, RFC5, MCM3, RAD51 |
| Cellular aldehyde metabolic process | 0.77 | 0.0123 | ALDH2, MGAT4A, ALDH1A1, IDH2, ALDH3A2, AKR1C3, ALDH7A1, IDH1, ALDH3A1 |
| DNA conformation change | 0.74 | 0.00056 | MCM5, GINS2, RPA1, BRIP1, CHTF18, MCM2, DSCC1, ERCC6L, RAD54L, RFC4, RFC5, TOP2A, MCM3, RAD51 |
| DNA-templated DNA replication | 0.73 | 3.37e-05 | CDC6, MCM5, GMNN, GINS2, RPA1, MCM2, RRM1, FEN1, PRIM1, ORC1, PCNA, RFC4, TONSL, RFC5, CCNE2, TIMELESS, MCM3, RAD51 |

DNA repair and replication.

Additional gene expression changes that may impact HUS pathogenesis include the IL10 signaling pathway that was upregulated in our 24-hour STX/LPS treatment and not at 4-hour treatment. A prior study demonstrated that IL10 deficient mice exhibit fewer neutrophils recruited to the kidney and increased survival in a HUS mouse model compared to control mice (28). LPS-induced inflammation may increase white blood cell delivery of STX to the kidney, contributing to renal EC STX-induced damage and HUS (4). Thus, LPS-induced IL10 secretion may contribute to LPS/STX synergistic toxicity by promoting neutrophil migration to the kidney and increased STX delivery to renal EC.

Although the STX-induced expression changes are delayed as compared to LPS effects in our HUVEC data, the synergistic differential gene expression changes may be responsible for the delay in TMA development. With both LPS and STX present during STEC infection, our data characterizing the synergistic EC expression changes with dual LPS/STX treatment provide a valuable resource for future examination of the pathophysiology of HUS and may represent a more accurate model of STEC induced differential gene expression than exposure to LPS or STX alone.

